# SMAC mimetics overcome apoptotic resistance in ovarian cancer through MSLN-TNF alpha axis

**DOI:** 10.1101/2024.01.24.576987

**Authors:** Ricardo Coelho, Brinton Seashore-Ludlow, Sarah Schütz, Flavio Christopher Lombardo, Elisabeth Moussaud-Lamodière, Ruben Casanova, Joanna Ficek-Pascual, Kathrin Brunhilde Labrosse, Michal Hensler, Monica Lopez-Nunez, Natalie Rimmer, Andre Fedier, Renata Lima, Céline Montavon Sartorius, Christian Kurzeder, Franziska Singer, Anne Bertolini, Tumor Profiler Consortium, Jitka Fucikova, Gunnar Rätsch, Bernd Bodenmiller, Olli Kallioniemi, Päivi Östling, Leonor David, Viola Heinzelmann-Schwarz, Francis Jacob

## Abstract

Resistance to chemotherapy and PARPi inhibitors remains a critical challenge in the treatment of epithelial ovarian cancer, mainly due to disabled apoptotic responses in tumor cells. Given mesothelin’s pivotal role in ovarian cancer and its restricted expression in healthy tissues, we conducted a drug-screening discovery analysis across a range of genetically modified cancer cells to unveil mesothelin’s therapeutic impact. We observed enhanced cell death in cancer cells with low mesothelin expression when exposed to a second mitochondria-derived activator of caspases (SMAC) mimetics, and demonstrated a compelling synergy when combined with chemotherapy in *ex vivo* patient-derived cultures and zebrafish tumor xenografts. Mechanistically, the addition of the SMAC mimetics drug birinapant to either carboplatin or paclitaxel triggered the activation of the Caspase 8-dependent apoptotic program facilitated by TNFLJ signaling. Multimodal analysis of neoadjuvant-treated patient samples further revealed an association between tumor-associated macrophages and the activation of TNFLJ-related pathways. Our proposed bimodal treatment shows promise in enhancing the clinical management of patients by harnessing the potential of SMAC mimetics alongside conventional chemotherapy.

## Introduction

Primary debulking surgery followed by platinum-based chemotherapy has been the cornerstone treatment for advanced high-grade serous ovarian cancer (HGSOC) (*1, 2*). In recent years, maintenance therapies including poly (ADP-ribose) polymerase inhibitors (PARPi), have become a valuable addition in patients with somatic or germline *BRCA1/2* tumors or homologous recombination deficiency (HRD), (*3–6*). Despite these therapeutic advancements, most patients with metastatic HGSOC inevitably experience disease recurrence and ultimately platinum-resistance disease (*7*). Considering these challenges, the necessity for overcoming drug resistance remains a critical clinical priority for these patients.

A growing body of evidence indicates that mesothelin (MSLN), a glycoprotein physiologically expressed at the surface of mesothelial cells lining the peritoneal cavity and overexpressed in several malignancies (*8–10*) plays a central role in tumor progression (*11–14*). Additionally, we identified MSLN as a key player in the multistep process of metastasis in EOC, where it promotes cell survival in suspension, invasion through the mesothelial cell layer, and peritoneal spread (*15*). Given its restricted expression and its role in ovarian cancer progression, MSLN has emerged as a promising candidate for targeted therapies using either antibody-based conjugates (*16, 17*) or CAR-T cell therapies (*18, 19*). Currently, several clinical trials (NCT02792114, NCT03816358, NCT04809766, NCT03814447, NCT03054298) are underway to assess the potential of these targeted therapies in combination with chemotherapy, an area that holds promise for preventing or overcoming drug resistance.

Despite the increasing promise of MSLN-targeted therapies in HGSOC, a notable gap exists in our understanding, particularly regarding the broader role of MSLN in overall drug responses, including its specific impact on chemotherapy responses. To fill this knowledge gap, we functionally address the question by conducting a comprehensive drug library screen on genetically manipulated cell lines with varying levels of MSLN expression. Furthermore, we identify drug synergies, specifically combining carboplatin or paclitaxel with SMAC mimetics (bimodal treatment) through a TNFa-caspase-8 dependent mechanism.

## Results

### Ovarian cancer cells with low MSLN expression are selectively targeted by SMAC mimetics

MSLN expression has been correlated with chemoresistance (*20,21*,*22*). However, these studies primarily report on associations without functionally elucidating the role of MSLN in drug responses. To fill this gap, we utilized our previously established experimental model (*15*), consisting of isogenic EOC cell lines with genetically manipulated MSLN expression, to a comprehensive drug sensitivity and resistant test (DSRT) comprising a 528 oncology drug library including kinase inhibitors, epigenetic modifiers, differentiating agents, and conventional chemotherapeutic compounds (<u>CHEMICAL COMPOUND LIBRARIES</u>, **Supplementary Table 1**). We evaluated cellular response to treatment by calculating the drug sensitivity score (DSS, (*23*)), which revealed a diverse spectrum across all cell lines tested (n=8). Despite marginal differences in DSS to conventional chemotherapeutics (**Figure 1A-B**), we identified SMAC mimetics as exhibiting a selective effect on MSLN^low/ΔMSLN^ EOC cell lines (**Figure 1C**). This increased sensitivity to SMAC mimetics was further validated for the SMAC mimetic drug birinapant in two EOC cell lines genetically engineered for MSLN expression, either by MSLN knockout (OVCAR3^Δ*MSLN*^) or MSLNoverexpression (OVCAR4 OE MSLN) (**Figure 1D**). To further prove the MSLN-specific response to SMAC mimetics, we conducted a co-culture time-lapse experiment exposing parental (MSLN^high^) and Δ*MSLN* OVCAR3 cells to birinapant treatment. For cell death assessment, we employed POPO™-1 iodide, a nontoxic cell impermeant dye. We observed a significant increase in cell death for Δ*MSLN* compared to parental OVCAR3, supporting the hypothesis that SMAC mimetics preferentially target MSLN^Δ*MSLN*^ cells (**Figure 1E**).

**Figure 1.**
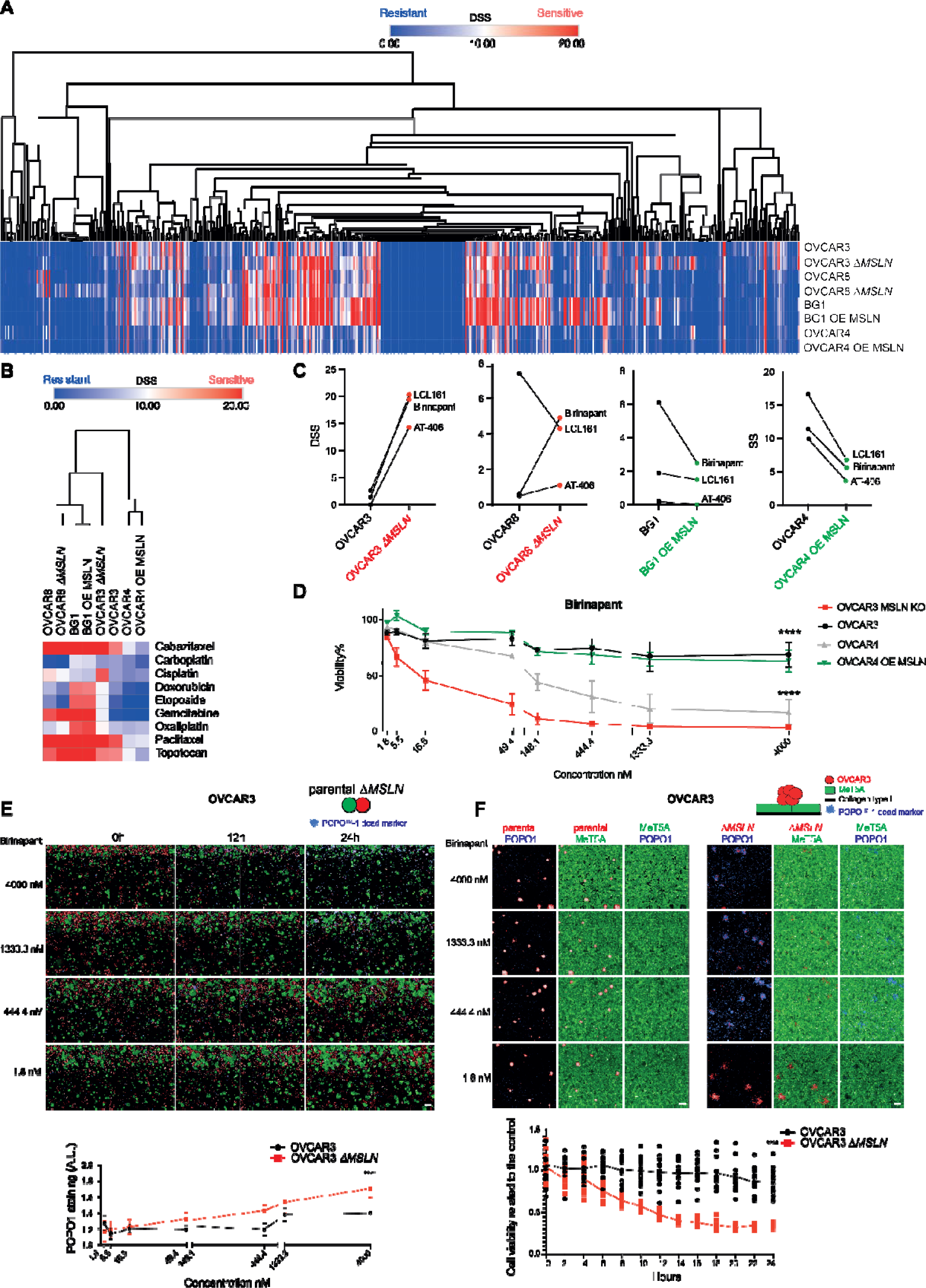
SMAC mimetics selectively target MSLN^low^ EOC cell lines. ***A.*** Heatmap depicting heterogeneous drug responses using a library of 528 compounds tested across 8 EOC cell lines with (Δ*MSLN for knockout; OE for overexpression*) and without genetic manipulation of MSLN expression. The represented values are drug sensitivity scores (DSS) colors indicating either sensitivity (red) or resistance (blue). **B.** Heatmap with drug responses of chemotherapeutic compounds commonly applied in EOC treatment. Visualization and hierarchical clustering of rows (1-Person correlation) was performed using Morpheus software. **C.** Dot plot illustrating the enhanced effectiveness of SMAC mimetics in cell lines characterized by low MSLN expression. **D**. Dose-response curves for birinapant across 4 different EOC cell lines. The data presented is the result of two independent experiments conducted in quadruplicate. **E.** Representative fluorescence images and quantification of a co-culture system using parental (EGFP) and Δ*MSLN* (dTomato) OVCAR3 cells treated with an increasing concentration of birinapant. The corresponding line chart summarizes the quantification of POPO™-1 iodide staining intensity after 48 hours of birinapant treatment. Scale bar 50µm. **F.** Representative fluorescent images of the mesothelial clearance assay in parental (left) and Δ*MSLN* (right) OVCAR3 cells with and without birinapant treatment. Cell viability was calculated based on POPO™-1 Iodide staining intensity and normalized against controls. Data is presented as mean ± SD of two independent experiments done in quadruplicate (D, E) or sextuplicates (F) *p* values were determined using two-way ANOVA followed by Sidak’s multiple comparison test. (*, *p* <0.05, ***, *p*<0.001, **** *p*<0.0001). Scale bar 50µm.

The tumor microenvironment (TME) plays a crucial role in modulating drug responses (*24*). Mesothelial cells represent a major cellular component of the peritoneal microenvironment, the location where EOC cells are predisposed to metastasize (*25–28*). Recognizing the importance of mesothelial cells in cancer progression and drug responses, we assessed the efficacy of birinapant in a co-culture assay using aggregates of parental and Δ*MSLN* OVCAR3 alongside normal immortalized mesothelial cells (MeT5A). This co-culture assay unveiled a selective effect of birinapant on Δ*MSLN* OVCAR3 cells, with minor effects on the MeT5A cell viability (**Figure 1F**). Taken together, our data indicates that the SMAC mimetics selectively targets MSLN^low/Δ*MSLN*^ EOC cells.

### SMAC mimetics restore chemotherapy sensitivity

Despite observing marginal differences in the cellular response to carboplatin and paclitaxel chemotherapeutics when comparing the DSS values among cell lines with differential MSLN expression, we hypothesized that chemotherapy might induce changes in MSLN levels. This hypothesis is based on the well-documented effect of chemotherapeutic agents in inducing cellular alterations, including DNA damage (*29*), and cell cycle arrest (*30*). These alterations ultimately lead to extensive modifications in gene expression and protein function (*31, 32*). Thus, we screened a panel of 18 EOC cell lines for MSLN expression and selected representative ones with MSLN^high^ expression for downstream assays (**Supplementary** Figure 1A). Following exposure to a gradient concentration of carboplatin and paclitaxel, the most effective chemotherapy compounds in EOC, we observed a decrease in MSLN protein levels across all cell lines tested (n=4). This reduction was accompanied by reduced cell viability, as evidenced by the increased expression of cleaved PARP (**Figure 2A, Supplementary** Figure 1B). The treatment of carboplatin combined with paclitaxel had a less pronounced impact on MSLN protein levels as if they were given as single compounds (**Supplementary** Figure 1C). Given this finding, we proceeded with downstream experiments using only the combination of SMAC mimetics with either carboplatin or paclitaxel. Of note, EOC cell lines used exhibited a general resistance to carboplatin and paclitaxel (**Supplementary** Figure 1D).

**Figure 2.**
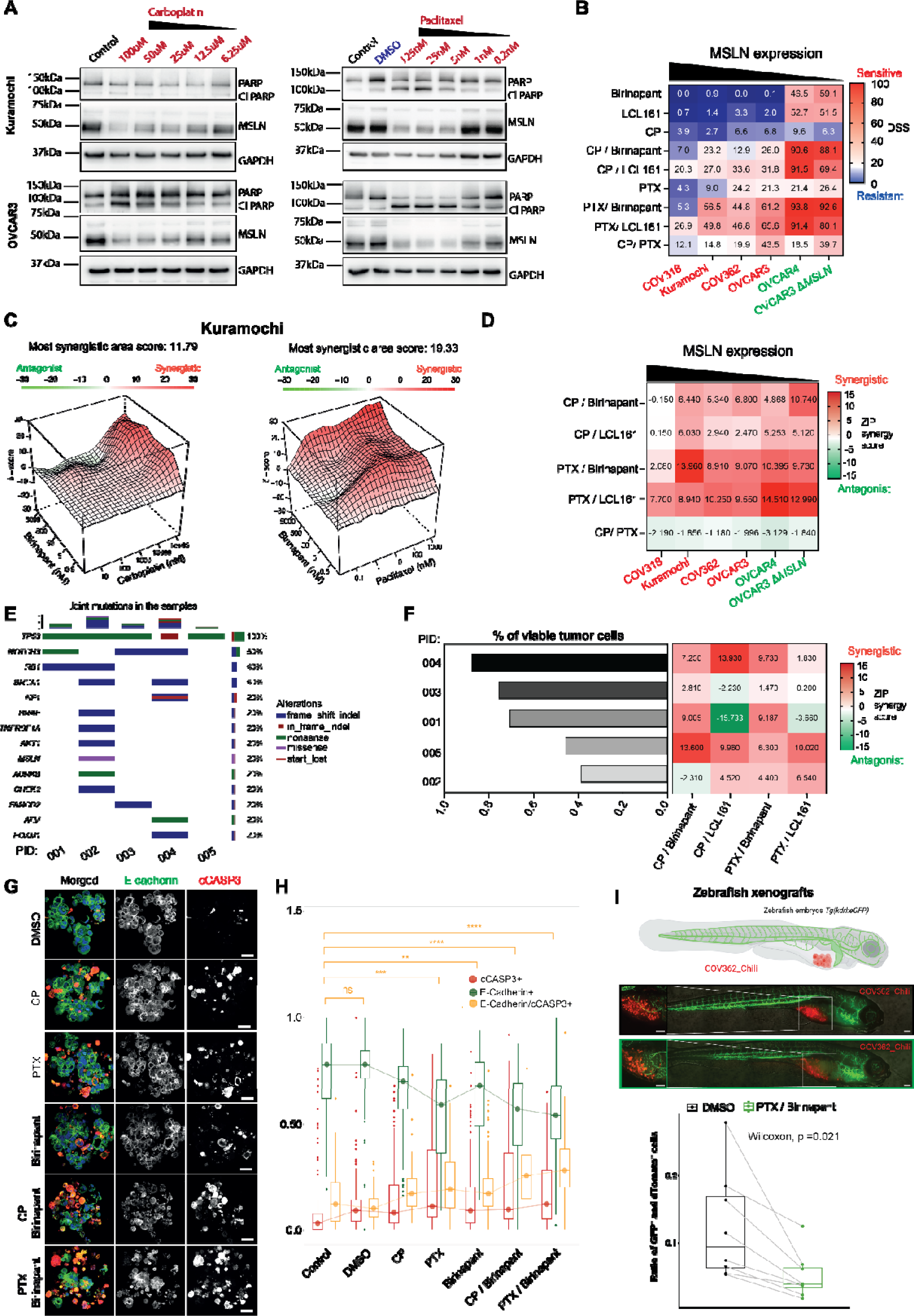
SMAC mimetics enhance chemotherapy sensitivity. **A.** Representative Western blots depicting reduction of MSLN following exposure to carboplatin and paclitaxel in Kuramochi and OVCAR3 cells. **B.** Heatmap with increasing DSS values for the combination of SMAC mimetics with either carboplatin or paclitaxel across six EOC cell lines, sorted by MSLN expression levels. **C** and **D.** Synergy scores for SMAC mimetics in combination with either carboplatin or paclitaxel. **E.** Oncoprint for HGSOC samples treated *ex vivo*. Matched genomic data obtained from whole-exome sequencing, only genes with alterations in at least one patient sample are visualized. **Supplementary Table 3**. **F.** Tumor content and matched synergy scores for SMAC mimetics in combination with either carboplatin or paclitaxel in HGSOC patient-derived cells. **G** and **H.** Representative immunofluorescent images and quantification of “on target” tumor effect of the different treatments using an *ex vivo* culture system in three HGSOC samples. Carboplatin, paclitaxel, and birinapant were used at the final concentration of 10µM, 10nM, and 500nM, respectively. At least 20 cell aggregates were imaged and quantified for each condition in each patient sample resulting in 122,805 cells analyzed. Scale bar 50 µm. *P* values were determined using the Wilcoxon comparison test *, *p* <0.05, ***, *p*<0.001, **** *p*<0.0001. **I.** Representative immunofluorescent images and quantification of COV362-Chili zebrafish xenografts, demonstrating a reduction in live COV362-Chili cells after 72 hours of exposure to the combination of birinapant (5000nM) with paclitaxel (1000nM). Data presented as a result of three independent experiments, each done in duplicate, *p* values were determined using the Wilcoxon comparison test. Scale bar 100µm.

Supported by the observed reduction in MSLN protein levels following exposure to either carboplatin or paclitaxel and the specificity of SMAC mimetics in targeting MSLN^low^ cancer cells, we conducted a dose-response matrix test and assessed the synergy scores for either carboplatin or paclitaxel in combination with SMAC mimetics (birinapant and LCL161) in a panel of EOC cell lines (n=6). We found enhanced MSLN-dependent drug efficiency when combining one chemotherapeutic compound (either carboplatin or paclitaxel) with an SMAC mimetic (bimodal treatment) (**Figure 2B**). To systematically analyze drug interaction patterns, we next employed the Zero Interaction Potency (ZIP) model (*33, 34*). This analysis revealed additive and synergistic effects for the bimodal treatments in six EOC cell lines tested in an MSLN-dependent manner. Surprisingly, the clinically routine combination of carboplatin with paclitaxel exhibited only a low interaction pattern compared to bimodal treatments. (**Figure 2C and D and Supplementary** Figure 2).

To further validate the synergistic effects, we profiled five HGSOC samples exhibiting high MSLN expression for *ex vivo* drug testing (**Supplementary** Figure 3A **and Supplementary Table 2**). These samples exhibited ubiquitous *TP53* mutations and heterogeneous mutation patterns in HR and other frequently altered HGSOC genes (**Figure 2E, Supplementary Table 3**). The dose-response matrix demonstrated a remarkable increase in the efficiency of SMAC mimetics in combination with either carboplatin or paclitaxel. We found additive and synergistic interactions of bimodal treatments for three out of five HGSOC patient-derived samples tested, importantly, independent of the tumor cell content (**Figure 2F and supplementary Figures 3B and C**). To assess the on-target tumor effect of the bimodal treatment, we next utilized a scaffold-free 3D *ex vivo* culture system followed by immunofluorescence staining for E-cadherin and cleaved caspase 3 (cCASP3). A significant increase in the percentage of apoptotic tumor cells was observed for the bimodal treatments as compared with controls suggesting a cancer cell-specific killing effect (**Figures 2G and H**). Cellular responses to our bimodal treatment were also assessed using the zebrafish embryo tumor xenograft model. In this model, we transplanted COV362 MSLN^high^ cells (carboplatin and paclitaxel-resistant) stably expressing dTomato fluorescent protein (COV362_Chili) into transgenic *Tg(kdrl:eGFP)* zebrafish embryos (*35, 36*). Zebrafish embryos were then exposed to birinapant in combination with paclitaxel or to the vehicle for 72 hours. The bimodal treatment significantly (p=0.021) reduced the percentage of viable COV362_Chili (DAPI^-^) cells as compared with the control condition (DMSO) (**Figure 2I)**. Additionally, to exclude any potential toxicity arising from off-target effects of the bimodal treatment, zebrafish embryos were exposed to a gradient concentration of paclitaxel, birinapant, and their combinations for 72 hours. Neither the mortality rate, malformations, missing organs, organ deficiencies, nor altered mobility capacity were observed in zebrafish larvae exposed to the different drugs and combinations, indicating low off-target toxicity of the tested compounds (**Supplementary** Figure 4). In summary, these results suggest that SMAC mimetics administered with either carboplatin or paclitaxel enhance tumor cell killing *in vitro*, *ex vivo*, and *in vivo* while causing minimal off-target toxicity.

### Birinapant promotes cancer cell apoptosis through TNF*α* signaling

SMAC mimetics have gained attention as a promising class of targeted therapies and are currently under clinical investigation for both solid and hematological malignancies (*37*). Mechanistically, SMAC mimetics are suggested to bind to Baculovirus Inhibitor apoptotic proteins Repeat domains (BIR) of Inhibitor of Apoptosis (IAP)s, activating NF-LJB signaling pathway and, induction of cell death in a TNFLJ-dependent manner (*38–40*). However, the detailed mechanism of action of SMAC mimetics, particularly regarding their synergistic effects when combined with either carboplatin or paclitaxel, remains unclear. To assess the impact of TNFLJ signaling on drug synergies, we exposed MSLN^high^ EOC cell lines (Kuramochi and OVCAR3) to the bimodal treatments in the presence of a neutralizing TNFLJ antibody or supplementing the culture medium with human recombinant TNFLJ protein. The results demonstrated that birinapant, used alone or in combination with either carboplatin or paclitaxel, significantly enhanced cancer cell death in a TNFLJ-dependent manner (**Figure 3A and B and Supplementary** Figure 5A-C). This was supported by the significant increase in the *TNFA* expression in EOC cells after exposure to carboplatin, paclitaxel, birinapant, and bimodal treatment (**Supplementary** Figure 5D). Of note, the major target of SMAC mimetics, cellular (c)IAP1 protein (*39*), exhibited a cell line-dependent protein expression after applied treatments (**Figure 3B and Supplementary** Figure 5C and E). These data suggest that birinapant may target different IAP proteins in a cell line-dependent manner, yet consistently produces an enhanced effect when combined with either carboplatin or paclitaxel.

**Figure 3.**
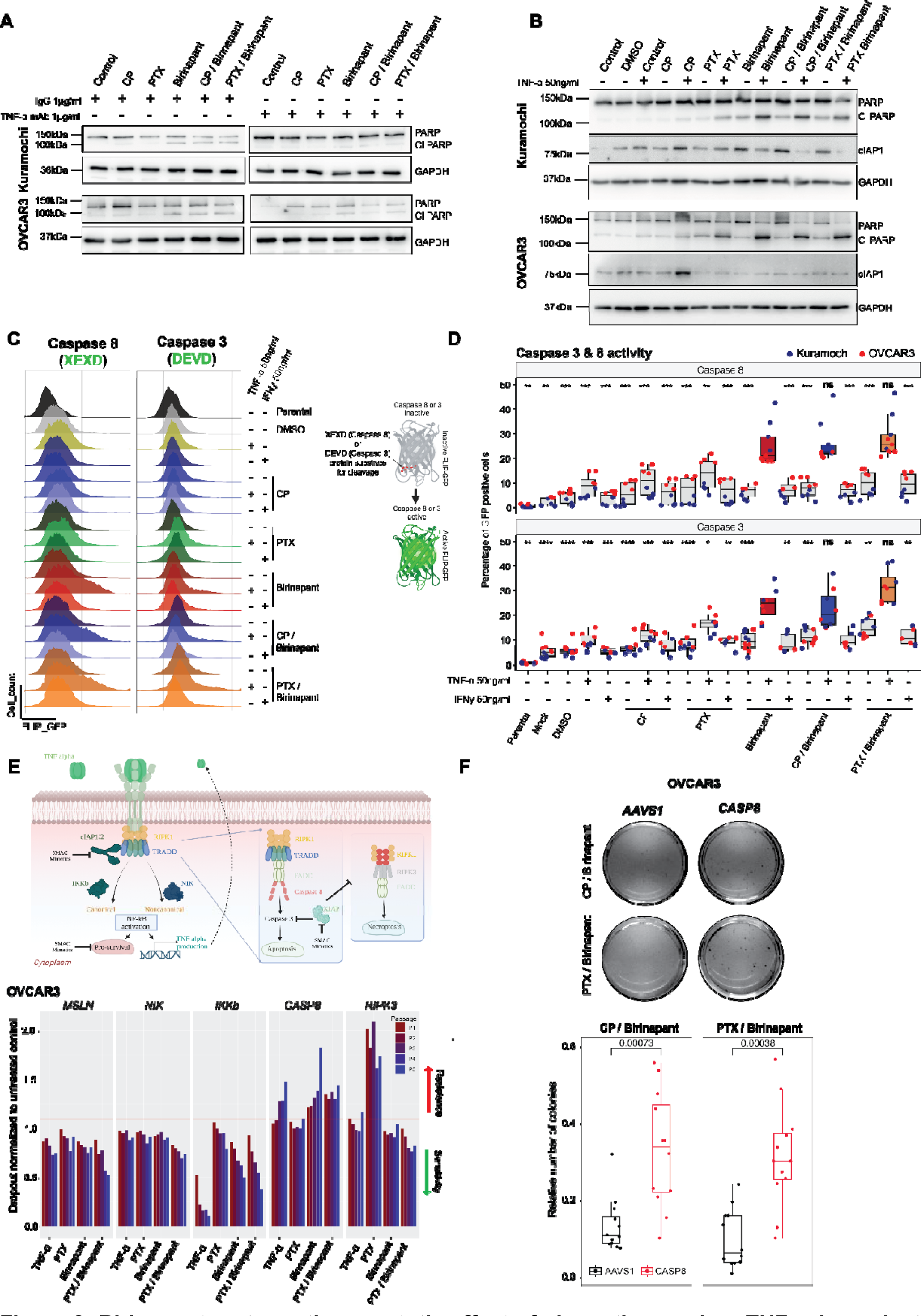
Birinapant restores the apoptotic effect of chemotherapy in a TNFL-dependen manner. **A** and **B.** Representative Western blots demonstrate the TNFLJ dependency for the enhanced efficacy of birinapant in combination with either carboplatin or paclitaxel in Kuramochi and OVCAR3 cell lines after 24 hours. Carboplatin, paclitaxel, and birinapant were used at the final concentration of 10µM, 10nM, and 500nM, respectively. **C** and **D**. Representative histograms and quantification of a real-time apoptosis reporter assay using FLIP-GFP as a readout supporting the specificity of TNFLJ in increasing cell apoptosis mediated through caspase 8 and 3 when combined with birinapant, and birinapant in combination with either carboplatin or paclitaxel. Carboplatin, paclitaxel, and birinapant were used at the final concentration of 10µM, 10nM, and 500nM, respectively. The data presented is the result of at least three independent experiments. *P* values were determined by Wilcoxon comparison, * *p* <0.05, ***, *p*<0.001, **** *p*<0.0001. **E.** Illustration of TNFLJ signaling pathway and histograms for CRISPR-Cas9 cell competition assay over five consecutive passages depicting increased resistance and sensitivity across different treatments upon gene editing of *MSLN* and TNFLJ pathways-related genes *NIK*, *IKKb, CASP8,* and *RIPK3***. F.** Colony formation assay with quantified colonies to assess treatment-dependent survival upon deletion of *AAVS1* (negative control) or *CASP8* in OVCAR3-Cas9^+^ cells. Carboplatin, paclitaxel, and birinapant were used at the final concentration of 1µM, 1nM, and 5nM, respectively. The data presented is the result of two independent experiments done in triplicates. *P* values were determined by Wilcoxon comparison test.

SMAC mimetics have been reported to trigger cell death through distinct programmed cell death (PCD) pathways, involving either caspase-dependent apoptosis or caspase-independent necroptosis (*41–45*). The induction of these different pathways of PCD depends on the specific combination of SMAC mimetics and the cancer model under investigation (*41, 43, 46, 47*). To evaluate which PCD is induced by our bimodal treatments in EOC, we established a reporter for Caspase 3 and 8 activity, leveraging a previously introduced FLIP-GFP system (*48*). Both reporter systems were validated by either immunofluorescence staining or pharmacological inhibition (Z-IETD-FMK) for Caspase 3 and 8, respectively (**Supplementary** Figure 6A and B). Flow cytometry data from Kuramochi and OVCAR3 cells, which stably expressed caspase reporters, demonstrated a significant increase in Caspase 8 and 3 activities (EGFP+) when TNFLJ supplementation was provided, either in combination with birinapant or as part of our bimodal treatments. Importantly, the increase in caspase activity was specifically induced by supplementation with TNFLJ and not by the IFNγ (**Figure 3C and D, Supplementary** Figure 6C).

To further validate apoptosis as the primary PCD pathway responsible for the observed synergies, and to investigate the role of the NF-LJB pathway in the outcome of bimodal treatment, we conducted a CRISPR-Cas9 cell competition assay (*49*) in OVCAR3 cells. In this assay, we targeted *NIK*, *IKKb*, *CASP8*, and *RIPK3* genes, encoding key players in the non-canonical and canonical NF-LJB pathway, apoptosis, and necroptosis, respectively. The generated data support the finding that apoptosis is indeed the primary PCD pathway triggered by the treatment with TNFLJ, birinapant, and birinapant in combination with paclitaxel. This was evidenced by the increased cell resistance to these treatments upon genetic deletion of Δ*CASP8* (**Figure 3E**). Enhanced sensitivity to the bimodal treatment-(birinapant/ paclitaxel) was also observed in Δ*KKb and* Δ*IKKb* indicating the additional involvement of the NF-LJB signaling pathway in the treatment response. The role of Caspase 8 in mediating cell death upon the bimodal treatment (birinapant with paclitaxel and birinapant with carboplatin) was further validated through clonogenic cell survival in Δ*CASP8* cells (**Figure 3F**) and by co-treatment with a Caspase 8-specific inhibitor (Z-IETD-FMK) (**Supplementary** Figure 6D). Paclitaxel monotherapy appeared to induce necroptotic cell death, as indicated by the increased resistance to paclitaxel observed in Δ*RIPK3* OVCAR3 cells (**Figure 3E**). Taken together, these results indicate that treatment of either paclitaxel or carboplatin simultaneously with birinapant primarily induces cell death through Caspase 8-mediated apoptosis. Furthermore, the results also suggest that activation of the NF-LJB pathway, whether through the non-canonical or canonical pathways, may modulate cellular responses to the applied treatment.

### TNF**LJ** pathway activation triggered by tumor-associated macrophages enhances cancer cell apoptosis

Despite initially responding to chemotherapy, 20% of HGSOC patients relapse within 6 months after completing primary chemotherapy treatment, while 60% of the patients relapse after 6 months (*50*). Patients who relapse after a treatment-free interval of more than six months often receive additional carboplatin or paclitaxel-based chemotherapy (*51*). However, multiple recurrences increase the likelihood of developing platinum resistance, consequently resulting in poor survival (*52, 53*).

To evaluate the potential of our bimodal treatment in enhancing chemotherapy effectiveness for patients with multiple relapses, we analyzed viable HGSOC cells derived from longitudinal samples collected at two disease recurrences (**Figure 4A**). Ascites taken at the time of the first and second relapse were subjected to *ex vivo* drug testing for carboplatin, birinapant, carboplatin with paclitaxel, and bimodal treatments. Overall, bimodal treatments revealed the highest efficacy in inducing tumor cell-specific apoptosis (E-Cadherin^+^, cCASP3^+^) compared to the remaining treatments tested (**Figure 4B** **and C**). Interestingly, bimodal treatments were generally more effective in reducing tumor cell viability at the second relapse time point compared to the first relapse.

**Figure 4.**
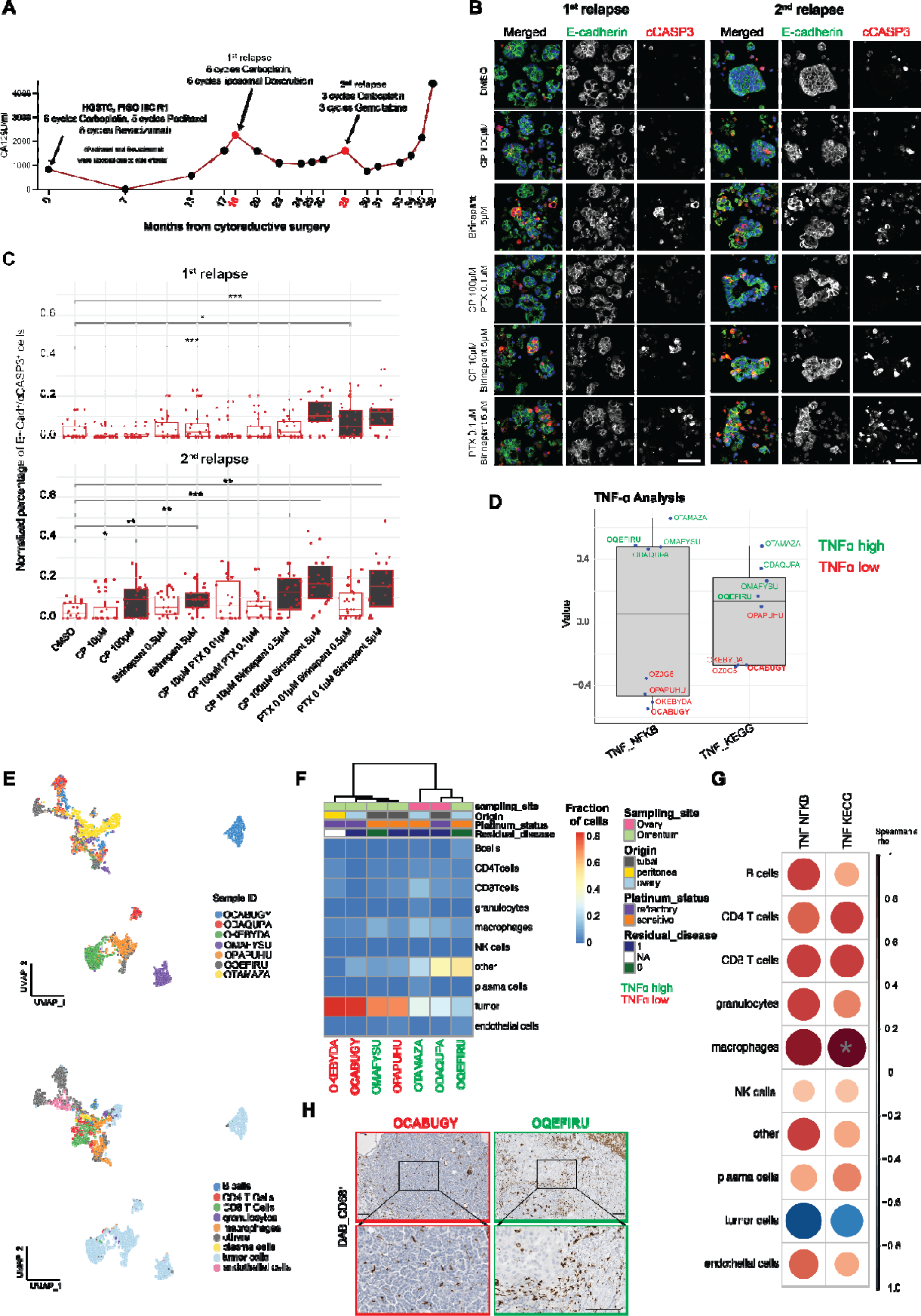
Macrophages in the TME promote TNF**LJ** pathway activation in HGSOC samples. **A.** Serum levels of CA125 in a patient with HGSOC during 36 months of follow-ups**. B** and **C.** Representative immunofluorescence images and box plots depicting increased cancer cell apoptosis (% of E-cadherin, cCASP3 positive cells) after exposure to birinapant in combination with either carboplatin or paclitaxel. At least 20 cell aggregates were imaged and quantified for each condition in each sample, in a total of 47’413 cells were analyzed. Scale bar 50µm. *P* values were determined by the Wilcoxon comparison test. \**p* <0.05, \*\**p* <0.01, \*\*\**p*<0.001. **D.** Pathway activation scores (z-scored) for neoadjuvant HGSOC-derived patient samples (n=8) with correction for the tumor content determined by CyTOF. Data generated from bulk RNAseq**. E.** UMAPs depicting the major cell type composition across seven different HGSOC samples using IMC. **F** and **G**. Heatmap and correlation matrix showing a significant association of macrophage abundance and TNFD -related pathways activation. A grey asterisk indicates significant correlations (r=0.82, *p*=0.034). **H.** Representative immunohistochemistry stainings in HGSOC samples with high and low CD68 cellular content. Scale bar 100µm.

In the complex ecosystem of tumor tissues, it is well-described that the tumor and the surrounding cells within the TME may be active participants in both the production and response to TNFLJ cytokine (*54, 55*). Considering the pivotal role of TNFLJ in the apoptosis-mediated drug synergies for our bimodal treatments, we performed an integrated analysis on samples derived from the Swiss Tumor Profiler (*56*). This analysis aimed to shed light on the potential contribution of different cell types within the TME to TNFLJ signaling activation and treatment response. This investigation included bulk RNA sequencing, cytometry by time-of-flight (CyTOF), and imaging mass cytometry (IMC) from the same sampling sites, obtained from patients with HGSOC who underwent neoadjuvant treatment. Based on our prior findings, we hypothesized that these patient samples were enriched in TNFLJ production post-neoadjuvant chemotherapy (**Supplementary** Figure 5D). Results from this analysis revealed a very strong positive correlation (spearman rho= 0.93, p=*0.007*, FDR=0.0317) between the TNFLJ gene sets and the fraction of tumor-associated macrophages (TAMs) (**Figure 4D-H**, **and Supplementary** Figure 7). To explore deeper into the IMC findings on the correlation between TAMs and TNFLJ gene sets, we conducted a deconvolution analysis of matching bulk RNA sequencing focusing on the immune compartment using CIBERSORT for relative quantification of the immune compartment using LM22 signature matrix (*57*). Our analysis revealed an enrichment of myeloid cells (monocytes and M2 macrophages) in samples exhibiting increased activation of TNFLJ gene sets (**Figure 5A**). Additionally, deconvolution analysis by applying the xCell score (*58*) and the Molecular Signatures Database (MSigDB) for Hallmarks signatures of TNFLJ signaling (https://www.pnas.org/doi/10.1073/pnas.0506580102) on bulk RNA sequencing data from an independent cohort of 97 HGSOC cases confirmed the enrichment of M2 macrophages and increased TNFLJ signaling activation in post-neoadjuvant-treated omentum metastasis tissue compared with chemotherapy-naive samples (**Figure 5B and Supplementary** Figure 8).

**Figure 5.**
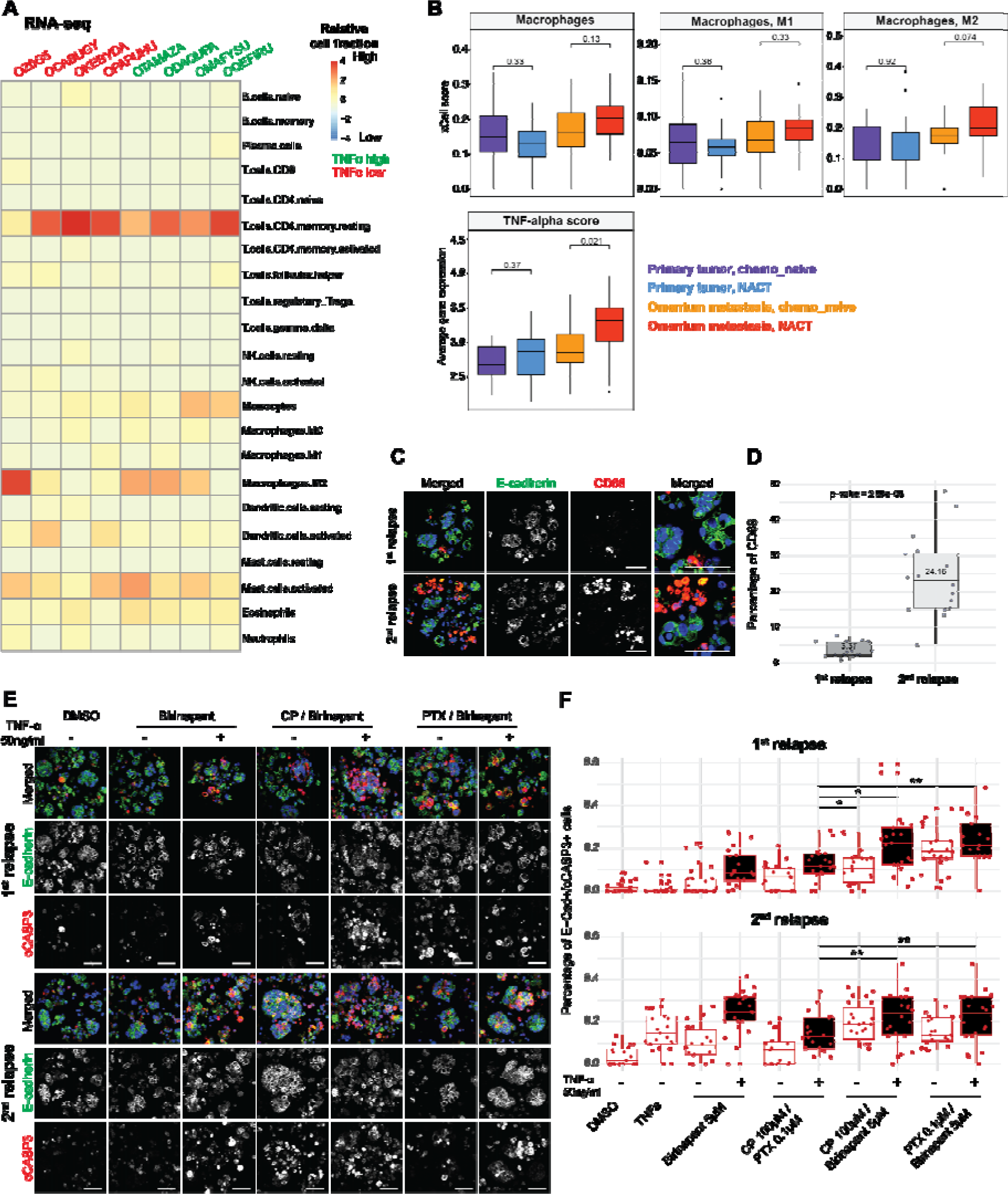
Macrophages and TNFL enhance response to bimodal treatments *ex vivo*. **A** Heatmap displaying the enrichment of myeloid cells (monocytes and M2 macrophages) in samples with higher (green) and low (red) TNFLJ gene set activation. Data sourced from bulk RNA sequencing, based on CIBERSORT relative quantification of the immune compartment using LM22 signature matrix. **B**. Boxplots depicting enrichment of M2 macrophages accompanied with increased TNFLJ pathway activation in omental metastasis in post-NACT samples. Data source from bulk RNA sequencing of 97 samples based on xCell score, Primary chemo_naive (n=30), Primary_NACT (n=24), Omentum metastasis_chemo_naive (n=19), and Omentum_metastasis_NACT (n=24). **C** and **D** Representative immunofluorescence images and corresponding box plots illustrate increased CD68^+^ cell percentage in the 2^nd^ relapse as compared with the 1^st^ relapse in PID_001. Quantification involved at least 16 cell aggregates that were imaged and analyzed for each condition in every sample, in a total of 3’354 cells analyzed. Scale bar: 50 µm, *P* values determined by *t*-test. **C** and **F** Representative immunofluorescence images and box plots showing increased cancer cell apoptosis (% of E-cad, cCASP3 positive cells) following bimodal treatments with TNFɑ supplementation as compared with carboplatin in combination with paclitaxel. Quantification included ≥ 20 cell aggregates that were imaged and quantified for each condition in each sample, in total 114’048 cells were analyzed. Scale bar: 50 µm. *P* values were determined by the Wilcoxon comparison test. \**p* <0.05, \*\**p* <0.01.

These findings prompted us to examine the impact of macrophage on responses to bimodal treatments in *ex vivo* cultures of longitudinal samples. Analysis of the percentage of macrophages (CD68^+^) across these samples revealed a higher CD68^+^ cell fraction in the second relapse, aligning with an enhanced cellular response to bimodal treatments compared to the first relapse (**Figure 4C** **and Figure 5C**). Additionally, supplementing *ex vivo* cultures with TNFLJ could further enhance the efficiency of the bimodal treatments (**Figure 5D and E**). In summary, these findings suggest a potential role for TAMs as a predictive biomarker for the response to bimodal treatments by regulating the TNFLJ pathway activation.

## Discussion

HGSOC is recognized as an extremely heterogeneous disease, presenting unique treatment challenges. Emerging therapeutic approaches aim to enhance patient outcomes by precisely identifying patient subgroups that will ultimately benefit from the applied treatment. The identification of such groups is based on multiple features of each tumor, including TME composition, genetic, and molecular profiles. Our approach seeks to improve patient stratification while maximizing the effectiveness of novel therapeutic approaches, thereby enhancing the quality of life and prolonging progression and disease-free survival. In this study, we introduce a novel approach to enhance the effectiveness of chemotherapy in HGSOC through co-treatment with SMAC mimetics. Our findings demonstrate that SMAC mimetics, particularly birinapant, in combination with either carboplatin or paclitaxel, trigger a Caspase 8-dependent apoptotic program facilitated by increased TNFLJ production and signaling activation. Through our analysis of neoadjuvant-treated HGSOC samples, we identified a positive correlation between TAMs and TNFLJ-related gene sets. Moreover, our experiments with HGSOC *ex vivo* cultures revealed a significant association between high TAM abundance and enhanced responses to the bimodal treatments. This suggests a potential role for TAMs in modulating drug responses through the mediation of TNFLJ signaling, highlighting their potential utility as predictive biomarkers for the proposed bimodal treatments.

Our study has uncovered a promising avenue in targeted therapy in combination with chemotherapy for HGSOC. Specifically, we have identified SMAC mimetics as a class of drugs that exhibit a preferential affinity for EOC cells exhibiting low MSLN expression. Additionally, our investigation has unveiled a compelling synergy between SMAC mimetics and standard chemotherapeutic agents with the potential to enhance the efficacy of treatment regimens in HGSOC patients. Notably, our results highlight the remarkable increases in TNFLJ expression following exposure to first-line chemotherapy. This, combined with validation experiments, indicates that TNFLJ acts as a driving force behind the observed synergies. Furthermore, we hypothesize that the reduction in MSLN protein levels in EOC cells, after exposure to carboplatin or paclitaxel chemotherapy, enhances cell death triggered by increased TNFLJ production in the presence of SMAC mimetics. This hypothesis is supported by a previous report showing increased resistance to TNFLJ-induced cell death in MSLN-expressing cancer cells (*11*).

While the exact mechanism through which MSLN contributes to heightened resistance to TNFLJ-induced cell death remains incomplete, previous studies have suggested that MSLN induces the activation of the NF-LJB pathway, resulting in the overexpression of multiple anti-apoptotic factors (*11, 59–61*), thereby contributing to increased resistance to TNFLJ-induced cell death (*11*). These findings align with data presented in the current study, where *MSLN* knockout cancer cells displayed enhanced sensitivity to TNFLJ, birinapant, and the combination of birinapant with paclitaxel. It is worth noting that, in contrast to TNFLJ, the supplementation with IFNγ did not result in increased activation of Caspase 3 and 8 after exposure to birinapant or bimodal treatments. However, previous studies reported that IFNγ supplementation increases caspase activity following exposure to SMAC mimetics (BV6) in breast and colon cell line cancer models (*46*). These differences between our results and the literature may suggest a cancer model-dependent effect of the combination of cytokines with SMAC mimetics, or dependent on the SMAC mimetic applied, considering their documented varying effects on target proteins (*39*). Additionally, as recently reported by Cremona *et al.*, the genetic background of cancer models can influence the response to SMAC mimetics. This influence was demonstrated recently by the enhanced response to SMAC mimetics in *BRCA1* mutant cells as compared with *BRCA* wild-type cells (*62*). Therefore, further investigation is necessary to elucidate the role of different inflammatory cytokines on the activation of different programs of PCD in combination with SMAC mimetics across various cancer models.

SMAC mimetics have been reported to activate both apoptosis and necroptosis, two distinct pathways of PCD (*43*). The pharmacological induction of necroptosis is considered a promising approach to overcome apoptosis evasion of tumor cells (*41, 63, 64*). However, necroptosis has a dual role in cancer, it can either promote or inhibit tumor growth, depending on the specific cancer type and context. As a fail-safe mechanism of PCD, necroptosis can prevent tumor growth in cases where apoptosis induction fails. Nonetheless, as a form of necrotic cell death, necroptosis can trigger inflammatory responses and reportedly promote cancer metastasis and immunosuppression (*65, 66*). In our present study, we demonstrated that the combination of the birinapant with either carboplatin or paclitaxel chemotherapy primarily induces cell death through Caspase-8-mediated apoptosis. These findings are inconsistent with a previous study that reported SMAC mimetics to enhance the antitumor effect of carboplatin combined with paclitaxel in ovarian cancer models through a Caspase 8-independent mechanism (*67*). In this work, the effect on cell death was reported in cells with Caspase 8 knockdown when a triple combination of birinapant, paclitaxel, and carboplatin was applied. However, when tested individually, birinapant exhibited synergy with paclitaxel but an antagonism effect with carboplatin in caspase 8-depleted cells (*67*). While we cannot dismiss that the inconsistencies between our results and the previous report might arise from different measurements and experimental setups, a possible explanation could relate to the use of a triple combination (birinapant, carboplatin, and paclitaxel). Of note, one of the two cell line models used, OVCAR8, has been reported to exhibit resistance to NF-LJB inhibition (*68*), indicating reduced responsiveness to TNFLJ signaling. Interestingly, Hernandez *et al.* also noted an increase in cell death for the combination of birinapant with either paclitaxel or carboplatin in Caspase 8 expressing models. While further investigations are necessary, these results imply marginal advantages in utilizing a triple combination of birinapant, carboplatin, and paclitaxel compared to the bimodal treatments.

The TME has been reported to influence tumor evolution by altering the milieu of growth factors and cytokines following chemotherapy (*69–71*). Changes in cytokine levels produced by stromal cells after chemotherapy tend to favor cancer cells expressing the corresponding receptors (*72*). In the current study, we found a compelling positive correlation between the presence of TAMs, specifically M2 like macrophages, and the activation of TNFLJ gene sets in neoadjuvant-treated HGSOC samples. Among the various cell types in the TME, macrophages are recognized as the major contributor to TNFLJ production and signaling activation (*73, 74*), which supports the findings of in the current study. Additional studies are required to fully understand the interaction and contribution of various cell types, especially TAMs, in the TME and their influence on TNFLJ signaling in tumor cells. This is particularly relevant in the context of the bimodal treatment where the synergistic effect of SMAC mimetics in combination with either carboplatin or paclitaxel depends on the TNFa signaling. By mechanistically unraveling the interplay between MSLN expression, TNFLJ signaling, TAMs, and the enhanced efficacy of bimodal treatments, we provide valuable insights that may guide the application of these therapeutic approaches in patients with recurrent HGSOC. While further research is needed to explore how different cell types in the TME contribute to TNFLJ signaling in tumor cells and the effects of combination therapies, our study established the groundwork for an exciting avenue in the treatment of this challenging disease. Specifically, our results suggest that TAMs enriched in the TME could potentially act as biomarkers for the proposed bimodal treatment, assisting in better patient stratification. Overall, our findings hold promise in improving patient outcomes, especially for those dealing with recurrent HGSOC.

## Material and methods

### Patient-derived cells

All HGSOC patient-derived samples and associated data included in this study fully adhere to ethical standards. Ethical approval was granted by the Ethical Committee of Northwest and Central, Switzerland (EKNZ, BASEC IDs: 2017-01900 and 2023-00988). All patients’ clinical characteristics are provided in **Supplementary Table 2**.

### Human cell lines

Ovarian cancer cell lines (Kuramochi, COV362, COV318, OVCAR3, OVCAR4, OVCAR5, OVCAR8, BG1, SKOV3, SKOV3 IP, ES-2, OVSAHO, A2780, A2780 cisplatin-resistant, TYKnu, TYKnu cisplatin-resistant, IGROV1, and IGROV1 paclitaxel-resistant) obtained from different commercial sources (**Supplementary Table 4**) were cultured in RPMI 1640 (Sigma-Aldrich) containing 10% fetal bovine serum (FBS) (Sigma-Aldrich) and 1% of penicillin/streptomycin (Sigma-Aldrich). Immortalized human mesothelial cell line MeT5A (ATCC) was maintained in Medium 199 (Sigma-Aldrich) containing 10% FBS, 3.3 nM epidermal growth factor (EGF) (PeproTech), 400 nM hydrocortisone (Sigma-Aldrich), 870 nM Bovine insulin (Sigma-Aldrich), 20 nM HEPES (Sigma-Aldrich) and 1:1000 of trace elements B (Corning). All cell lines were maintained at 37°C and 5% CO_2_ authenticated using short tandem repeat (STR) profiling (Microsynth AG) and regularly tested for the absence of mycoplasma (*75*). To study the role of MSLN in drug sensitivity and resistance we used a previously established and characterized cell line models: parental (MSLN^high^) and MSLN knockout (Δ*MSLN*) OVCAR8 and OVCAR3 cell lines, as well as parental (MSLN^low^) and MSLN overexpressing (MSLN OE-MSLN^high^) BG1 and OVCAR4 cells (*15*). These EOC cell lines were initially selected based on their nature of MSLN expressing levels.

### Drug sensitivity and resistance testing (DSRT)

We used the DSRT system developed at the Institute for Molecular Medicine Finland (FIMM) for drug library screening. The FIMM oncology library-FO (https://www.helsinki.fi/en/infrastructures/drug-discovery-chemical-biology-and-screening/infrastructures/high-throughput-biomedicine/chemical-compound-libraries), consists of 528 approved and investigational oncology drugs including target compounds such as kinase inhibitors, epigenetic modifiers, differentiating agents and conventional chemotherapeutics. The compounds were dispensed in 384-well plates (Corning) at five different concentrations covering 1 and 10000nM range, using acoustic liquid dispensing technology (Echo 550; Labcyte Inc.). Assay-ready plates were stored in pressurized StoragePods (Roylan Developments) under an inert atmosphere until used. Before DSRT, growth patterns, and rates were evaluated by viability assays to ensure logarithmic cell growth during the 72 hours of drug exposure. Using a MultiDrop^TM^ Combi dispenser (Thermo Fisher Scientific) 5μL of culture media (RPMI 1640 supplemented with 10% FBS and 1% Pen/strep) was dispensed into assay-ready plates followed by a brief centrifugation. Next, ovarian cancer cells (OVCAR3, OVCAR8, BG1, and OVCAR4) and mesothelial cells, MeT5A, were seeded at the density of 1.5 to 2x10^3^ cells/well in 20μl of media using the Multidrop Combi dispenser (Thermo Fisher Scientific) into drug-containing plates followed by brief centrifugation and incubated for 72 hours at 37°C in the presence of 5% CO_2_. Cell viability/cellular ATP levels were measured with CellTiter-Glo 2.0 (CTG, Promega) with detection on the Ensight plate reader (PerkinElmer). Raw data were further processed to calculate the drug efficacies of individual drugs. Using in-house software, data from the plate reader were normalized per plate to the percent of cell viability in control wells (DMSO, and benzethonium chloride-BzCl). Concentration-response curves were fitted to percent viability values using a four-parameter logistic model and processed further to a sensitivity metric (drug sensitivity score; DSS) using a weighted area-under-the-curve calculation (*76*). For the validation experiments, hit compounds and controls (DMSO, 100 μM BzCl) were pre-dispensed on 384-well plates (Corning) using the Echo 550 (Labcyte Inc). Each compound was plated as quadruplicate in eight concentrations (3-fold dilution).

### Dose-response matrix analysis and synergy scoring

For the synergy experiments, we used 384-well screening plates with five concentrations per compound in duplicate (carboplatin in combination with paclitaxel) or triplicate (birinapant, LCL161 in combination either with carboplatin or paclitaxel, all dissolved in the same solution). Cell seeding, viability, and calculation of DSS values (for cell lines and patient-derived samples, **Supplementary Table 2**) were performed as described above for DSRT. Response matrices were then calculated using the ZIP reference model with the SynergyFinder web application (version 3.0; synergyfinder.fimm.fi) (*34*). Combinations with a score > 10 were considered to exhibit a strong synergy and <-10 a strong antagonism. Scores between 5 and 10 were considered moderately synergistic, between -5 to -10 as moderately antagonistic, and between -5 and 5 were indicative of additive effects.

### Evaluation of drug efficiency using live cell confocal microscopy

Parental and Δ*MSLN* OVCAR3 cells were manually seeded into 384-well CellCarrier culture plates (PerkinElmer) at the density of 2x10^3^ cells/well in 20μl of media (RPMI 1640 supplemented with 10% FBS and 1% Pen/strep) and incubated for 24 hours. For the co-culture assay, equal amounts of each cell population (1000 cells/well) were seeded in 20μl of media. After the incubation time, 5μl of POPO™-1 Iodide (Thermo Scientific) diluted in culture media (final concentration of 0.2μM) was added to all the wells of the plate. Birinapant (4000nM to 1.8nM concentration) was added in 5μl of media to the wells in quadruplicate as well as a negative (DMSO 0.1%) and positive control (BzCl 100μM) followed by time-lapse microscopy using the OperaPhenix microscope (PerkinElmer) for 48 hours. Image acquisition and quantification of POPO™-1 iodide staining intensity were done using the Harmony high-content imaging and analysis software 4.8 (PerkinElmer). Mesothelial clearance was performed by seeding MeT5A-GFP cells (*15*) at the density of 1.3x 10^4^ cells/well into 384-well CellCarrier plates (PerkinElmer), pre-coated with 10μg/ml collagen type I (Sigma-Aldrich), in 20μl of media. In parallel, ovarian cancer cell aggregates were generated by seeding the cells into t25 micro cavity flasks (Corning) followed by a 24-hour incubation. After incubating, aggregates (10μl ∼7 aggregates) were transferred to the plate containing the MeT5A-EGFP cells and exposed to a gradient concentration of birinapant (4000nM to 1.8nM). POPO™-1 Iodide death marker (Thermo Fisher Scientific) was added to all the wells of the plate at the final concentration of 0.2μM followed by 24-hour time-lapse microscopy using the OperaPhenix microscope (PerkinElmer). Image acquisition and quantification of POPO™-1 Iodide (Thermo Fisher Scientific) staining intensity was done using the Harmony high-content imaging and analysis software 4.8 (PerkinElmer).

### Immunoblotting

EOC cell lines were lysed in 1x radioimmunoprecipitation assay buffer (RIPA, Cell Signaling Technology, BioConcept) containing a proteinase inhibitor cocktail (Sigma-Aldrich). Lysates were clarified by centrifugation at 18,000 g for 15 minutes at 4°C. Clarified lysates were quantified using the Pierce TM BCA protein assay (Promega) and boiled in 1x sample buffer (50 mM Tris-HCl, 1% SDS, 100 mM DTT, and 10% glycerol) at 95°C for 5 min and resolved by SDS-PAGE. Proteins were then transferred to a polyvinylidene difluoride (PVDF) membrane (BioRad) and blocked with 5% (w/v) bovine serum albumin in TBS-T (20 mM Tris-Base, 150 mM NaCl, pH 7.8, 0.1% Tween 20) for 1 hour at room temperature. The membrane was incubated with one of the listed primary antibodies (**Supplementary Table 5**) diluted in 5 % (w/v) BSA in TBS-T overnight at 4°C. After washing (3 times, 10 minutes) in TBS-T, membranes were incubated with corresponding HRP-conjugated secondary antibodies (1:10000, Cell Signaling, BioConcept) for 3 hours at room temperature. Finally, the membrane was incubated with Super Signal West Dura Extended Duration Substrate (Thermo Fisher Scientific) for the detection of the HRP signal. Western blot results were visualized by Gel Doc XR+TM (BioRad) and analyzed by Image Lab^TM^ software (BioRad). Original membrane images can be found in the **Supplementary** Figure 9-13.

### *Ex vivo* patient-derived cultures

Tissue or ascites samples were thawed (retrospective cohort), and digested into single cells (tissues) using a mix of collagenase/dispase (Roche), accutase (Millipore), DNAse type IV (New England Biolabs/ Bioconcept) in DMEM/F12 (Sigma-Aldrich) for 1 hour at 37°C under agitation. After digestion, samples were filtered through a 70µm cell strainer and washed with PBS (Sigma-Aldrich). Cells were then seeded into agarose chips, which were pre-made using MicroTissues 3D petri dish micro-mold (Sigma-Aldrich), at a density of 2.5x10^5^ cells/ chip in 200µl of Ovarian TumorMACS^TM^ (Miltenyi Biotec) medium and incubated for 24 hours. Patient-derived cells were then exposed to the different drugs and their combinations for 48 hours followed by fixation with 4% formalin (Formafix AG) for 20 minutes at room temperature. After fixation cells were washed with PBS, and to prevent cell loss in the subsequent steps, a thin layer of liquid histogel (Epredia) was added to the top of the cells. Agarose chips were then subjected to a standard paraffin embedding protocol and sectioned at 5µm thickness followed by immunofluorescence staining.

### Immunofluorescence

After deparaffinization, heat-induced (98°C) antigen retrieval was performed with a citrate buffer (pH 6.0) (Thermo Fisher Scientific), and slides were incubated with Triton X-100 0.25% in PBS for 5 minutes, washed in PBS for another 5 minutes followed by incubation with blocking solution (5% FBS with 1% BSA in PBS 1% Triton-X 100) for 1 hour. After blocking, samples were incubated with one of the listed primary antibodies (**Supplementary Table 5**) diluted in 1% BSA 0.1% Triton-X 100 in PBS and incubated overnight at 4°C. Primary antibodies were detected using goat anti-mouse or anti-rabbit IgG (H+L) Alexa fluor 488 or 647 (Cell signaling Technologies/ Bioconcept) secondary antibodies diluted 1:500 in 1% BSA 0.1% Triton-X 100 in PBS for 3 hours in the dark at room temperature. Slides were mounted using ProLong® Gold antifade reagent (Cell Signaling Technology, BioConcept) and a coverslip. Images were taken using the confocal microscope Nikon CSU-W1, analyzed with Image J (2.3.0/1.53q) and Qupath (0.3.0) software, and developed scripts for cell detection and annotations. Further analyses were performed by R/Bioconductor.

### Zebrafish embryo tumor xenograft

All zebrafish experiments and husbandry followed federal guidelines complied with the Swiss Animal Protection Ordinance and were approved by the Cantonal Veterinary Commission of Basel-Stadt. The xenotransplantation experiments were performed as previously described with minor alterations (*35, 77*). Zebrafish embryos were anesthetized in 0.02% tricaine (Sigma-Aldrich) at 2-day post fertilization (dpf) and 250–350 COV362 cells (stable expressing dTomato fluorescent protein) were microinjected into the vessel-free area of the yolk of a transgenic Tg(*kdrl:eGFP*) line. Embryos harboring cells were incubated at 35°C in 1x E3 medium containing either DMSO 0.1% or birinapant in combination with paclitaxel at the final concentration of 5000nM and 1000nM, respectively. Three days after transplantation, zebrafish larvae (8-9 per condition) were dissociated into single cells by applying enzymatic digestion using 5µl of liberase (Sigma-Aldrich) (2.5mg/ml) in a final volume of 50µl in PBS and incubated for 15 minutes at 32°C with agitation 800 rpm. This incubation step was repeated three times with repeated vigorous pipetting in between resulting in a total digestion time of 60 minutes. After complete digestion, Liberase was inactivated by adding 5µl of 5% BSA in PBS. An additional 150 µl of PBS was added, followed by filtration through a 70µm filter. The filter was rinsed once with PBS to collect all cells. Single cells were washed twice with PBS and then stained with a DAPI (Biolegend) solution (0.2 µg/ml) and analyzed on a CytoFLEX S (Beckman Coulter) for DAPI-negative and dTomato-positive cells. The normalization for the input was performed by dividing the number of dTomato^+^ cells by the number of eGFP^+^ (zebrafish) cells. Toxicity tests of the various compounds and combinations were performed in zebrafish embryos with a modified protocol based on the previously described methods (*78, 79*). Zebrafish embryos (at 2 dpf) were incubated for 3 days at 35°C in 1x E3 medium containing a gradient concentration of DMSO, paclitaxel, birinapant, and birinapant in combination with paclitaxel. After the incubation period, the mortality rate and general behavior were macroscopically evaluated (n=2 wells per condition, each with 5 larvae). At least two larvae per condition were selected for microscopic assessment of the morphology using confocal microscopy (Nikon CSU-W1). This evaluation included a detailed analysis of malformation, absence of organs, or signs of organ dysfunction.

### Lentivirus production and transduction

Lentivirus production was performed using the HEK29T cells that were cultured in RPMI 1640 with 10% FBS and 1% penicillin/streptomycin. One day before transfection 4x10^6^ cells were seeded into a 75cm^2^ tissue culture flask. For each flask, 4µg of the plasmid encoding the targeted information, pUltra (Addgene #24129), pUltra-Chili (Addgene #48687), LentiV-Cas9-puro (Addgene #108100), LGR2.1 (Addgene #108098) encoding the gRNA targeting the gene of interest, pLenti-CMV-Blast-FLIP-GFP-DEVD, and pLenti-CMV-Blast-FLIP-GFP-XEXD, and 2µg of packaging (pCMVR8.74, Addgene #22036) and envelope (pMD2.G Addgene #12259) plasmids were transfected using 24µl of jetPEI reagent and 1 ml of 150mM NaCl solution (Polyplus-transfection). Culture media was changed 24 hours after transfection. The supernatant containing active lentivirus particles was collected 48 hours later, filtered with a 0.45µm filter (Sartorius), and stored at −80°C. Target cell lines were then transduced with the desired lentiviral supernatant. If needed after 3 passages, cells were enriched for dTomato, EGFP, or mCherry positive cells using the BD FACS Aria Cell sorter (BD Bioscience).

### Generation of caspase 3 and caspase 8 reporters

Plasmid encoding the sequence of the cleavage site for Caspase 3 (DEVD) and FLIP-GFP reporter (Addgene# 124428) was digested using the *NheI*_HF and *XmaI* (New England Biolabs, Bioconcept) and cloned into the pLenti_CMV_GFP_Blast (Addgene #17445) backbone, that has been previously digested with *XbaI* (New England Biolabs, Bioconcept) and *XmaI*, followed by transformation into *Stbl3 E. coli* strain. The right insertion and sequence of FLIP_GFP_DEVD in the pLenti_CMV_GFP_Blast backbone (pLenti_CMV_GFP_Blast_FLIP_GFP_DEVD) was confirmed by Sanger DNA sequence (Microsynth) using the CMV-F primer (**Supplementary table 6**). Caspase 8 reporter was established by replacing the DEVD amino acid sequence with XEXD in pLenti_CMV_GFP_Blast_FLIP_GFP_DEVD plasmid through the In-fusion Snap assembly master mix (Takara Bio) using the following oligos forward: 5’-GTCTGGAAACCGATGGTGGCGGTGGCAAGGTG-3’ and reverse 5-CATCGGTTTCCAGACCTGATGCATCGGTAATGCCAG-3’, PCR reaction was conducted under the following conditions: denaturation at 98°C for 10 seconds, followed annealing at 55°C for 5 seconds and 72°C for 50 seconds, repeated for a total of 35 cycles. The resulting PCR product was then purified using the Macherey-Nagel NucleoSpin gel and PCR clean-up kit (Thermo Fisher Scientific). Linearized vector (67.4ng) was then mixed with 5X In-Fusion Snap Assembly Master Mix and nuclease-free water in a final volume of 10ul. This mixture was incubated at 50°C for 15 minutes followed by transformation into Stellar™ competent cells a *HST08 E.coli* strain. The protein-encoding XEXD DNA sequence was confirmed by the Sanger DNA sequence using the CMV_F primer (**Supplementary Table 6**).

### Caspase 3 and Caspase 8 activity analysis

Caspase 3 and Caspase 8 activity was visualized by flow cytometry in BG1, COV362, COV318, OVCAR4, OVCAR3, and OVCAR8 cells stably expressing the FLIP-GFP reporter for caspase 3 and 8 activity. Cells (5x10^4^/well) were seeded into 48-well plates and incubated for 24 hours, followed by exposure to carboplatin, paclitaxel, birinapant, TNFLJ, IFNy, and co-treatments for 24 hours. Cells were then harvested and stained with DAPI (0.1µg/ml) solution. Caspase 3 and 8 activity (EGFP^+^ cells) was analyzed using the CytoFLEX Flow Cytometer (Beckman Coulter) and Flowjo v10 BD (Becton Dickinson) software. Caspase 3 and 8 activity was represented as the percentage of EGFP^+^ cells within the mCherry positive cells (cells with stable expression of the caspase 3 or caspase 8 reporter). For each cell line, two to four independent experiments were performed.

### Molecular cloning for the single RNAs

Single guide RNAs (sgRNA) targeting protein-coding DNA sequences of target genes was designed using Benchling (Biology Software, 2021, retrieved from https://benchling.com).

SgRNAs with high-quality scores were selected for editing of the target gene (**Supplementary Table 7**). Single-strand oligonucleotides were purchased from Sigma-Aldrich annealed and cloned into LRG2.1 (addgene, #108098) using the *BsmBI* endonuclease restriction site and T4_DNA ligase (Promega) for subsequent expression of the sgRNA together with EGFP fluorescent protein. Ligations were transformed into *Stbl3 E.coli* following ampicillin selection, ZR Plasmid Miniprep Classic plasmid purification (ZYMO Research, Lucerna-Chem), and Sanger DNA sequencing (Microsynth) to confirm insertion of respective sgRNA using the human U6 primer (**Supplementary Table 6**).

### CRISPR-Cas9 cell competition assay

CRISPR-Cas9 cell competition assay was performed as previously described (*49*). Stable Cas9-expressing ovarian cancer cells were transduced with lentivirus particles containing sgRNAs targeting selected genes (**Supplementary Table 7**). SgRNAs targeting the same gene were pooled and further diluted in a 1:5 or 1:8 ratio with a complete culture growth medium. The medium was changed 24 hours after transduction and cells were incubated for 2 additional days until measuring the EGFP-positive cells. The percentage of fluorescence-positive cells was determined initially (baseline reference, passage 0, 3 days after transduction) and all following passages using the CytoFLEX Flow Cytometer (Beckman Coulter). In each passage, cells were washed with FACS-wash (FW-1% FBS in PBS) in 96-well V-shape plates and stained with DAPI (0.1 µg/ml) (Biolegend) for 5 minutes. All investigated cell lines were gated individually to exclude debris, doublets, and dead (DAPI^+^) cells.

Delivery of compounds and combinations was initiated at passage 1 and cells were kept under treatment until passage 5 (20-23 days after transduction). The following concentrations were used: 50 ng/ml, 1-2uM, 1-2nM, and 2-5nM of TNFLJ, carboplatin, paclitaxel, and birinapant. Dropout values represent the fold-change of the percentage of EGFP^+^ cells at each passage, relative to passage 0, and were used as a readout of effects on cell fitness or drug sensitivity conferred by the CRISPR-*Cas9* mediated gene mutations. Data analysis was performed using FlowJo v10 BD (Becton Dickinson) and R/Bioconductor scripts. Relative drug sensitivity was calculated by dividing the dropout values for the control condition (targeted gene without treatment) by the values for the target gene under treatment.

Of note, due to the high number of guides applied in this study, each assay was performed at least in two independent experiments with the negative control *AAVS1* (non-essential gene) and at least one positive control (*RPA3* or *PCNA*). The robustness of the competition assay is described in (*49*).

### Colony formation assay

Drug sensitivity was determined by the colony formation assays. OVCAR3-Cas9 cells were transduced with lentivirus supernatant encoding sgRNAs targeting *AAVS1* or *CASP8*. Three passages after lentivirus transduction cells were enriched for EGFP^+^ cells using the BD FACSAria Cell sorter (BD Bioscience). Cells were then seeded into 12 well plates at a density of 600 cells/well and exposed to the different drug combinations at the final concentration of 1µM, 1nM, and 5nM for carboplatin, paclitaxel, and birinapant, respectively. Complete growth media with or without drugs was replaced every 3 days. After two weeks, colonies were fixed at room temperature for 1 hour and stained with 0.05% crystal violet (Sigma-Aldrich) in 4% formalin (Formafix AG) the plates were then rinsed 3-4 times with water, dried, and images of the well were taken using the Fusion FX7 Edge Imaging System (Witek AG).

### RT-qPCR

To extract RNA, 3x10^5^ cells were seeded in 12-well plates and incubated for 48 hours. OVCAR3 and Kuramochi cells were then exposed to the following compounds and combinations for 24 hours: carboplatin (10µM), paclitaxel (10nM), birinapant (500nM), and birinapant (500nM) combined with either carboplatin (10µM) or paclitaxel (10µM). Total RNA extraction was performed using the RNeasy Plus universal mini kit (Qiagen). RNA was eluted in 30µl of RNase-free water and the concentration was measured using a NanoDrop (Thermo Fisher Scientific). A total of 1µg of RNA in 20µl reaction volume was reverse transcribed using the iScript Reverse Transcription Supermix for RT-PCR (Bio-Rad). RT-qPCR was performed for *TNFA*, and reference genes *SDHA* and *HSPCB* (*80*) in a 10µl reaction containing 25ng cDNA (initial total RNA), 400nM forward and reverse primer (**Supplementary Table 6**), nuclease-free water, and 1x GoTaq Master mix with ROX as a reference dye (Promega) on a Thermo ABI 7500 fast real-time PCR machine (Applied Biosystem, Thermo Fisher Scientific). Quantitative PCR was performed in triplicates and analyzed using the 2–ΔΔCt method.

### TNF**LJ** supplementation and neutralization

OVCAR3 and Kuramochi cells were seeded at a density of 5x10^5^ cells/ well in a 6-well plate and incubated for 48 hours. Cells were then exposed for 24 hours to the following compounds and combinations: carboplatin, paclitaxel, birinapant, carboplatin and birinapant, paclitaxel and birinapant either supplemented with TNFLJ (50ng/ml, Peprotech) or with TNFLJ neutralizing antibody (1µg/ml, Cell Signalling Technologies), following protein extraction and immunoblot analysis. Rabbit IgG (Santa Cruz Technologies) was used as a non-specific binding control.

### Whole exome sequencing

Exonic regions were captured using the Twist Biosciences Human Comprehensive Exome Panel for pair-end 100bp reads, as described by the manufacturer, and sequenced on an Illumina NovaSeq 6000. Post-sequencing analysis of Whole Exome Sequencing (WES) genomic data included the following steps: Initially, adapter trimming was performed using SeqPurge (v2022_04), retaining reads with a minimum length of 15 bp. Alignment to the reference genome GRCh38 was executed with BWA MEM (v0.7.17), followed by post-processing via Samtools (v1.11) and Picard (v2.27.1) for removal of PCR duplicates and secondary alignments. Base recalibration was carried out using the GATK suite (v4.2.6.1). Quality control metrics were assessed using the following tools: FastQC (v0.11.9), Picard, GATK, Samtools, and QualiMap (v2.2.2d), with MultiQC (v1.9) for aggregating the QC results. Variant calling was performed with a combination VarScan2 (v2.4.4), Mutect2 (GATK v4.2.6.1), and Strelka2 (2.9.10), with variants reported by at least two of three variant callers retained for downstream analyses. Variants from all samples were merged using BCFTools (v1.15.1) and annotated using the Ensembl Variant Effect Predictor (VEP) (v108) with databases including ClinVar, dbSNP, gnomAD, dbNSFP, and COSMIC.

### Suspension and imaging mass cytometry

Cytometry by the time of flight and imaging mass cytometry experiments (CyTOF and IMC respectively) were performed as described before (*81*). For CyTOF, single cells were identified using a random forest classifier trained on a subset of manually gated cells using the Cytobank platform (*82*). For IMC, cell types were identified using a series of cell clustering and random forest classification steps.

### Sequencing, alignment, and counting

Bulk RNA libraries were ribo-depleted and sequenced with the Illumina NovaSeq 6000 system. Alignment to the reference genome (GRCh38) was performed with STAR (*83*). The raw expression counts were obtained using simple counting, an in-house pipeline accessible through <u>github:gromics</u> (at commit #67e1f4e).

### Sample quality control

Quality control of samples was performed based on their RNA Integrity Score (RIN) and the following FASTQC metrics: “Per Sequence GC Content”, “Overexpressed Sequences”, “Per Base Sequence Quality”, “Per Sequence Quality Scores” and “Per Base N Content”. If multiple FASTQ files were generated per sample, e.g., due to sequencing separately forward and reverse reads or sequencing in multiple lanes or flowcells, the FASTQC metrics were aggregated per sample, with at least one FAIL/WARN resulting in FAIL/WARN for this metric for the considered sample. To determine the overall sample QC, taking into account all listed FASTQC metrics as well as RIN score, the following criteria were adopted: RIN below six and three or more failed FASTQC modules resulted in classifying a sample as FAIL, RIN score above 7, and less than two failed FASTQC modules classified samples as PASS, anything in-between or with missing information categorized sample as WARN. Applying these criteria to the cohort resulted in 3 x FAIL, 11 x WARN, and 38 x PASS, with both WARN and PASS samples being considered for preprocessing and all downstream analyses.

### Data preprocessing

Only protein-coding genes, according to Gencode release v32 (GRCh38.p13) were kept, except for mitochondrial genes, and further filtered to a set of genes with more or equal to 10 counts in the cohort (sum across all samples). For pathway analysis, three subsequent preprocessing steps were taken: a variance stabilizing transformation (VST), batch-effect correction, and regressing out effects of confounding variables. VST, which internally adjusts for differences in library sizes between samples, was performed with DESeq2 (*84*), without taking any design information into account. Moreover, batch correction with remove BatchEffect from the limma package (*85*) was carried out using a sequencing date as a batch (with a total of 6 batches). Furthermore, the first principal component capturing differences in RIN score (Pearson’s r: - 0.68), as well as tumor content (estimated using CyTOF data) were regressed using the remove Batch Effect function.

### Pathway activation scoring

The pathway definitions, in terms of gene lists, were obtained from MSigDB (<u>link</u>) and KEGG (<u>link</u>) databases, accessed on 29.07.2023. Pathway activation scores were computed using the Gene Set Variation Analysis (GSVA, (*86*) with the minimum gene set size per pathway set to five, Gaussian kernel, and using the normalized enrichment score (mx.diff=TRUE). The pathway activation scores were then z-scored across the whole cohort, before subsetting the data to neo-adjuvant cases only.

### Bulk RNA sequencing

RNA-seq data was generated using the Illumina TruSeq Stranded mRNA Library Prep with 100 paired-end 100bp reads, according to the manufacturer’s protocol, and sequenced on an Illumina NovaSeq 6000. The analytical pipeline was built around the nf-core RNAseq pipeline (<u>v3.8.1)</u>. In brief, FASTQ files underwent trimming and quality control using Trim Galore (v0.6.7), and Cutadapt (v.3.4) followed by mapping to the GRCh38 primary assembly using STAR (v 2.7.10a), and the Gencode v41 primary annotation. Subsequent quantification was performed with Salmon (v1.5.2). Quality control (QC) metrics were evaluated using featureCounts (subread v2.0.1) for biotype counts, DESeq2 (v1.28.0) for Principal Component Analysis (PCA) and sample similarity, Picard MarkDuplicates (v1.28.0) for identifying and marking duplicate reads, Samtools (v1.15.1) for various alignment and post-alignment metrics, FastQC (v0.11.9) for basic quality metrics, and RseQC (v3.0.1) for assessing RNA-seq data quality. The QC results were collated into a singular report using MultiQC (v1.11). RNA-seq data were normalized using the variance stabilizing transformation of the DESeq2 R (v1.36.0) package to correct for technical variances. On this data, the deconvolution analysis was performed by using a Docker container pulled locally to perform matrix deconvolution using the LM22 signature matrix, allowing for the quantification of relative abundances in immune cells, based on the bulk expression matrix of about 600 genes provided by the LM22 file.

## Statistical analysis

Statistical analysis and figures were obtained using the software R Studio version 3.6.1 and 4.2 both versions were used interchangeably (www.R-project.org) and the {tidyverse} R software suite. All negative and positive controls were performed on multiple replicates at least 3 times in each cell line. All experiments were performed at least in duplicate and statistical evaluation was performed using Prism 9 software (https://www.graphpad.com/scientific-software/prism/) or R/Bioconductor. Where applicable, evaluation was done using two-way ANOVA with correction for multiple comparison tests using BH (Benjamini-Hochberg correction) or the Wilcoxon test. *p*-values of <0.05 were considered statistically significant and presented as a value or as \*\**p*LJ<LJ0.01, \*\*\**p*LJ<LJ0.001, \*\*\*\**p*LJ<LJ0.0001.

## Acknowledgments

We are grateful to the Flow Cytometry Core Facility (Stella Stefanova and former members), Microscopy Facility (Mike Abanto, Loïc Sauteur, and Ewelina Bartoszek), Histology Facility (Diego Calabrese), Martina Konantz, Markis Affolter group (Biozentrum, University Hospital Basel, Basel, Switzerland) for their assistance and support during the zebrafish experiments, and to the TuPro consortium and Miltenyi Biotec for providing essential support and reagents.

## Author contributions

Conceptualization: RCo, FJ, LD, and BS-L

Methodology: RCo, BS-L, NR, FCL, and FJ

Investigation: RCo, SS, FCL, RL, MNL, NR, JF, RC, KBL, FS, AB, JF, MH and ELML

Data curation: RCo, FCL, and FJ Writing—original draft preparation: RCo and FJ

Writing—review, and editing: all authors read and approved the final manuscript

Visualization: RCo, FJ, FCL, RC, and JF

Supervision: LD, PO, OK, BS-L, VHS, and FJ

Project administration: LD, VHS, and FJ.

Funding acquisition: RCo, LD, VHS.

